# What is Known, Unknown, and Needed to be Known about Damage Caused by Wild Pigs

**DOI:** 10.1101/2023.10.27.564386

**Authors:** Kurt C. VerCauteren, Kim M. Pepin, Seth M. Cook, Sophie McKee, Abigail Pagels, Keely J. Kohen, Ingrid A. Messer, Michael P. Glow, Nathan P. Snow

## Abstract

Damage assessments provide evidence for initiating and evaluating management programs that protect natural resources and human livelihoods against invasive species. Wild pigs (*Sus scrofa*) cause extensive damage in their native and non-native ranges, but the extent of current damage and efficacy of management of the damage (i.e., population control of wild pigs, exclusion fencing, etc.) remains poorly described. We conducted a systematic review of physical damage caused by wild pigs to summarize what is known and identify knowledge gaps for damage assessment. Wild pig damage assessments have been overwhelmingly qualitative (84% of studies) and measured differently across studies, which prevents the determination of typical damage amounts to a particular resource and comparison across studies. Key priorities going forward are to standardize damage assessments quantitatively and measure population density (or index of such) of wild pigs concurrently with damage assessments to determine the relationship between damage and population levels. We provide a framework for inferring damage in new areas and assessing the benefits of management – to evaluate and optimize landscape-scale management programs. Overall, we recommend future studies strive for: 1) report the amount of damages in a standardized fashion (e.g., area damaged/area surveyed), 2) evaluate and report the amount of damage relative to the density of wild pigs, and 3) when reporting economic costs of damages incurred and management actions, describe the economic valuation method used along with the year of reference for the valuation. Capturing these elements are necessary steps to predict the benefits of management for an area with particular profile of resources and wild pig density, even in areas where damage assessments are not available. Meeting these criteria with allow for more generalizable results that can inform managers across the nearly global distribution of wild pigs

## Introduction

Wild pigs (*Sus scrofa,* called wild boar in their native range and wild or feral pigs, hogs, or swine in their non-native range) were once restricted to Eurasia and North Africa, but are now present on all continents except Antarctica (Barrios-Garcia and Ballari 2012). Introduced wild pigs are a varying mixture of domestic pigs, Asiatic wild boar, and European wild boar (Wehr et al. 2018). Wild pigs wreak havoc on ecosystems, agriculture, and personal property around the globe in both their native and invasive ranges (Barrios-Garcia and Ballari 2012; Bengsen et al. 2014; Bevins et al. 2014; Massei and Genov 2004). They can affect ecosystem function and biodiversity values (Bengsen et al. 2014; Wehr et al. 2018), cause agricultural damage (Anderson et al. 2016, Anderson et al. 2019; McKee et al. 2020), and transmit disease to humans, livestock, and wildlife (Barrios-Garcia and Ballari 2012; Bevins et al. 2014). Estimated costs of those damages and associated efforts to control populations of wild pigs are very high (e.g., $1.5 billion annually in the USA: Pimentel 2007). While we know that these damages occur globally, there is still much we do not know about damage caused by wild pigs and how it can be effectively managed (Nogueira et al. 2009; Barrios-Garcia and Ballari 2012; Bengsen et al. 2014; Wehr et al. 2018).

The full scope of damage from any pest or invasive species that is expanding geographically has two components: 1) damage that is occurring to specific resources (current damage), and 2) damage that will occur where species are likely to expand or increase populations (potential damage). Information from these current or potential damages shape the establishment and regional objectives of management programs. For example, a program may strive ‘to reduce current damage by some amount’ or ‘to prevent additional damage to specific resources or other regions’. Damage information is important for evaluating program benefits, which may fuel decisions about future funding, and be used to identify management strategies that optimize management objectives (Adams et al. 2005; Bengsen et al. 2014; Bevins et al. 2014; Ballari et al. 2015).

Additionally, although many stakeholders view wild pigs negatively, at least in their non-native range, some maintain high societal and cultural significance of wild pigs (Nogueira et al. 2009). The impacts of wild pigs can be categorized as positive (i.e. ecosystem services), negative (i.e. damage), or neutral depending on a range of factors such as the observer (e.g., farmer, hunter, etc.), socio-cultural values (e.g., indigenous, rural, etc), and perceived value of affected resources (e.g., seed consumption, seed dispersal, personal injury, etc). In this review, we focused primarily on the negative damages and assessments of those damages, but also acknowledged the neutral and positive impacts of wild pigs. Comprehensive damage assessments can help all stakeholders make more informed decisions about the costs and benefits of different management decisions and what level of management is acceptable, which can improve management outcomes (Enck et al. 2006). Thus, damage assessment is a critical tool both before and during management programs (Massei and Genov 2004; Bengsen et al. 2014).

We conducted a comprehensive review of studies that documented physical damage by wild pigs around the globe to determine the state of knowledge on wild pig damage, identify gaps with current methodologies, and identify priorities for future damage assessment work. We review why and how damage has been measured, what has been learned about damage, what remains to be learned, and identify methodological gaps. For this review, we considered types of damage from wild pigs as: rooting, trampling, wallowing, and consumption of native and agricultural plants and animals (Gray et al. 2020), competition with native species for resources, vehicle collisions, personal property damage, and in rare cases, direct attacks on humans (Mayer and Brisbin 2009). We summarized economic losses when reported, but we focused on reviewing the non-monetary amounts of physical damage and how to measure and manage that damage. This review builds from the recent review by Didero et al. (2023) on economic estimates of damage from wild pigs by expanding outside of the USA, and by focusing on a more inclusive list of studies reviewed. We did not limit our studies to just those that reported an economic value for damage, nor did we restrict types of physical damage. We also captured information on management and population monitoring techniques, which has not been a focus previously.

We ended our effort by presenting a conceptual framework demonstrating how damage assessment studies are the cornerstones for development and evaluation of management programs, especially when multiple studies can be collated for application to the same objective (i.e., program evaluation or comprehensive damage assessment). We identify research priority for damage assessments to fulfill this purpose effectively. Finally, we make recommendations for future studies of damage caused by overabundant pest species.

## Methods

We performed a systematic review of all studies with physical damage (or related impacts – positive or negative) and wild pigs in their native and nonnative range. Our criteria for inclusion were that the study measured direct damage (e.g. wild pigs rooting of row crops) or indirect damage (e.g., effects of wild pigs rooting on butterfly communities by altering vegetation; Scandurra et al. 2016) caused by wild pigs. We included damage to crops, pasture, livestock, natural resources (e.g., vegetation, water, soil properties, wildlife), sensitive species, public recreation (e.g., golf courses, hunting opportunity), historical sites, and personal property. We excluded disease studies from our damage categories because most addressed different objectives (not damage assessment) and thus used different methods. We excluded review articles from our database because we were interested in analyzing how primary research studies were measuring damage, however we used review articles to help locate primary literature.

We queried Web of Science in August 2023 using the following statement in the ‘topic’ field: (boar OR pig OR hog OR pigs OR scrofa) AND (feral OR wild OR invasi*) AND (damag* OR depredat* OR rooting OR wallow* OR graz* OR digging OR disturbance OR “ecological impact” OR predate* OR vehicle OR collision). This generated 1498 articles. We also conducted a series of similar queries in Digital Commons to gather grey literature that may not be published in Web of Science.The total unique number of publications from the Digital Commons searches was 202. Finally, when we found relevant review articles on wild pig damage, we searched the reference lists to extract papers that met our criteria but did not show up in our Web of Science or Digital Commons searches. Of the 1700 papers we found using both search engines, 122 studies from the Web of Science and five studies from Digital Commons met our criteria. We also identified 82 studies in 15 review articles and book chapters, for a total of 209 studies that met our criteria. Four articles (Choquenot et al. 1997; Tierney and Cushman 2006; Vilardell et al. 2008; Higginbotham and Bodenchuk 2014) had two separate studies that we treated as such and thus there were 205 unique papers but 209 studies.

We developed a database with fields describing the source of the article (how it was located, title, author, year), the objective of the study, information about what type of damage was studied, how it was studied, what value of damage was recorded, what type of management if any occurred, and what the wild pigs density was if it was measured (Table S1, Text S1). Each paper was reviewed by two separate reviewers to cross-validate the database. When different entries were found we reexamined the paper to come to a consensus on the entry. All numerical entries in the database were standardized to appropriate metric units and currency was converted to United States Dollars (USD) 2021 using an inflation calculator (https://www.usinflationcalculator.com/; USD values before 2021 were inflated to present day values; values reported in Euros were converted to USD using https://www.google.com/finance/quote/EUR-USD before conversion to 2021 USD).

## Why damage is measured

The 209 damage assessment studies (Table S1, Text S1) we reviewed had a variety of objectives (Table 1, Figure 1). General trends were that a large majority of studies included qualitative damage assessment (i.e., whether or not damage occurred, or whether the effect seemed stronger on one resource relative to another) as the only objective (Table 1), but that there is an increasing trend for obtaining quantitative estimates (i.e., numerical measures of how much damage) and considering multiple quantitative objectives simultaneously. For example, the emphasis on qualitative damage assessment decreased slightly during the period that damage assessment studies were increasing exponentially (2001-2021; Figure 1) relative to the earlier period (1970-2000; Figure 1). In contrast, quantitative damage assessment studies only accounted for 38% of all damage assessment study objectives, but increased over time (Table 1). Objectives of determining optimal management strategies or assessing attitudes towards wild pigs and their damage were infrequent but showed the greatest proportional gain over time (Table 1, Figure 1). Studies that included joint objectives of quantifying damage, evaluating impacts, and/or determining optimal management strategies also increased proportionally over time.

**Figure 1.** Counts of studies by decade since 1970 that include different study objectives. We categorized the objectives as: 1) to conduct qualitative assessments where the objective was to assess if damage occurred, to determine its effect size, or to determine environmental or biotic factors that led to more severe damage, 2) to conduct quantitative assessments where the main objective was to measure the amount of damage in monetary or non-monetary terms, 3) to assess the impacts of management on damage (i.e., evaluation), including if or by how much damage was reduced and the identification of management strategies that reduce damage, 4) to determine the benefits of controlling damage by different management strategies, and 5) to determine stakeholder attitudes towards wild pigs and/or management of their damage

**Table 1.** Focus of studies during the lower interest time frame (1970-2000) and higher interest time frame (2001-2021).

| **Study objective** | **1970-2000**<br>**Count (%)** | **2001-2021**<br>**Count (%)** | **Overall**<br>**Count (%)** |
| --- | --- | --- | --- |
| Qualitative damage assessment | 43 (96%) | 132 (80%) | 175 (84%) |
| Quantitative damage assessment | 13 (29%) | 67 (41%) | 80 (38%) |
| Evaluation of management impacts | 10 (22%) | 46 (28%) | 56 (27%) |
| Determining optimal management strategies | 1 (2%) | 9 (5%) | 10 (5%) |
| Measuring stakeholder attitudes | 1 (2%) | 14 (9%) | 15 (7%) |
| Quantitative damage<br>assessment & evaluation<br>of management impact | 7 (15%) | 30 (18%) | 37 (18%) |

## What is known about damage caused by wild pigs

Wild pigs affect resources negatively both in and outside their native range of Eurasia and North Africa (Figures 2 and 3). In their non-native range, damage to natural resources (i.e., ‘Ecological Impacts’) are most widely studied, whereas in the native range damage to agricultural crops are most widely studied (Figure 3). In their non-native range the same amount of research interest has been invested in studying damage to crops as in studying damage to personal property. Rooting is the was most commonly reported (Figure 4) with more than 63% of studies reporting some form of rooting damage. Damages from vehicular collisions and human injury accounted for 12.0% of studies reported. Damage from wallowing or trampling were the least reported at 10% each. These findings indicate that wild pig foraging behavior accounts for the majority of the damage reported, especially to agriculture and natural resources. This is not surprising considering these damages also generate the largest economic consequences (Didero et al. 2023). Although most studies report wild pig effects as negative impacts, there have been positive or neutral effects reported, especially to natural resources (Figure 3).

**Figure 2.**
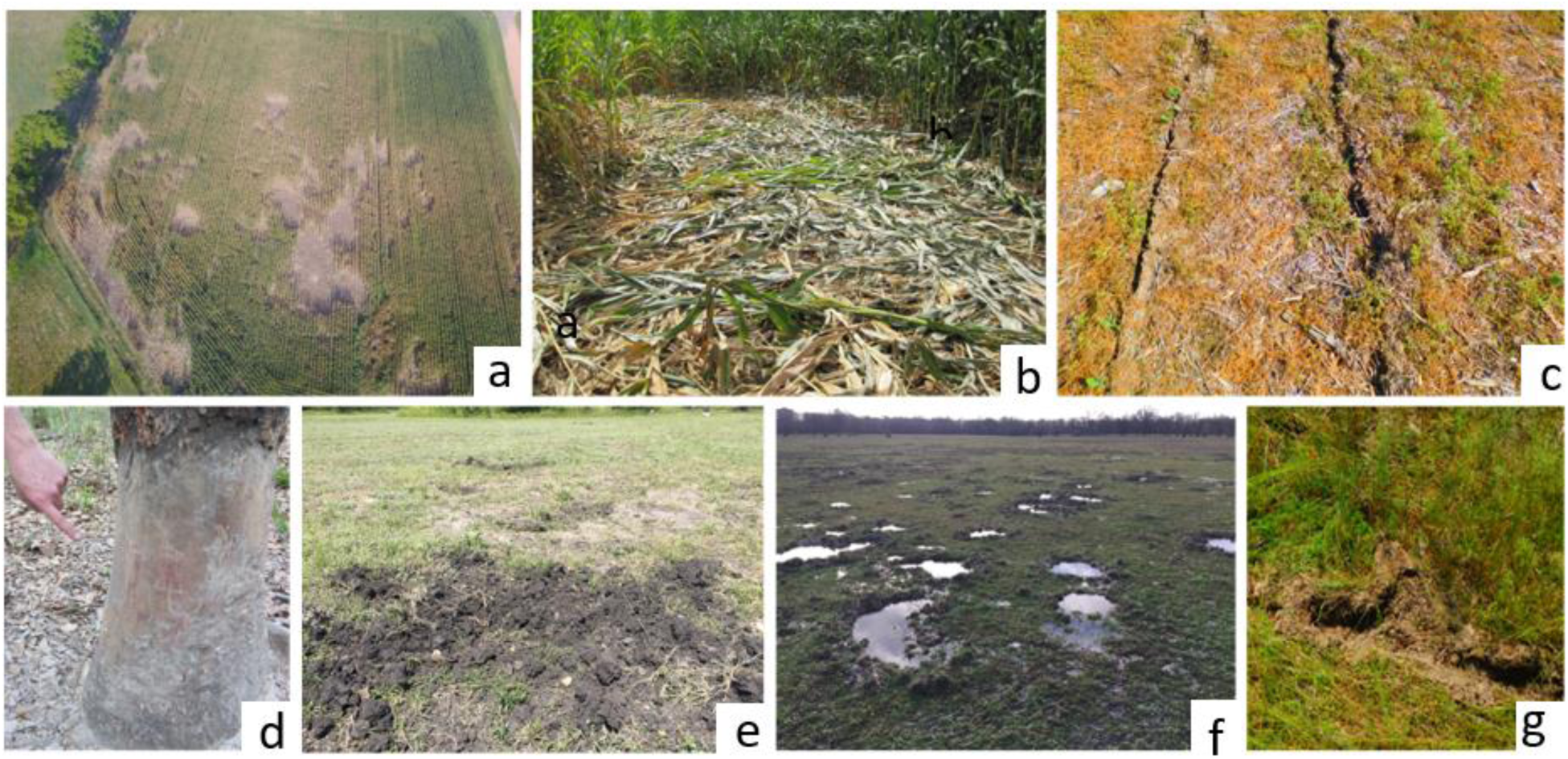
Examples of damage cause by wild pigs. a: Aerial view of damage to a corn field, b: Ground view of damage to corn, c: Rooting to consume a row of planted seeds, d: Debarking and girdling of a tree, e: Pasture rooting, f: Wetland impacts from wallowing, g: Pasture damage from wallowing

**Figure 3.**
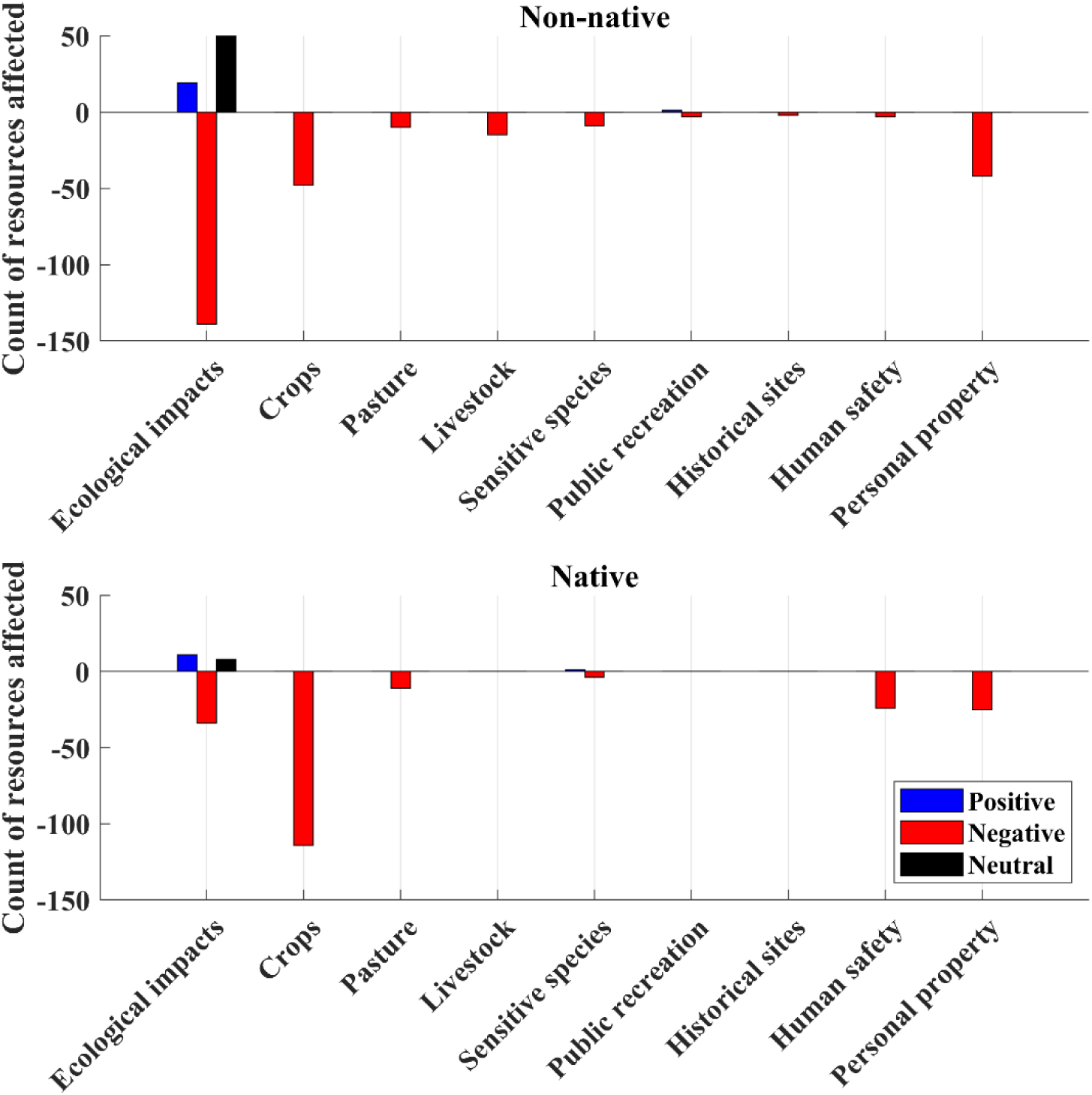
Effects of non-native (top) and native (bottom) ranges of wild pigs on different resources. Bars indicate a count of the number of resources across all papers that were impacted negatively (red), positively (blue), or unaffected (black) by wild pigs. Some of the 209 papers included an assessment of effects on multiple resources (e.g., crop types, soil properties, plant species, etc.) and thus the count is more than the number of papers. ‘Ecological impacts’ includes natural resources (vegetation, wildlife), soil components, and water

**Figure 4.**
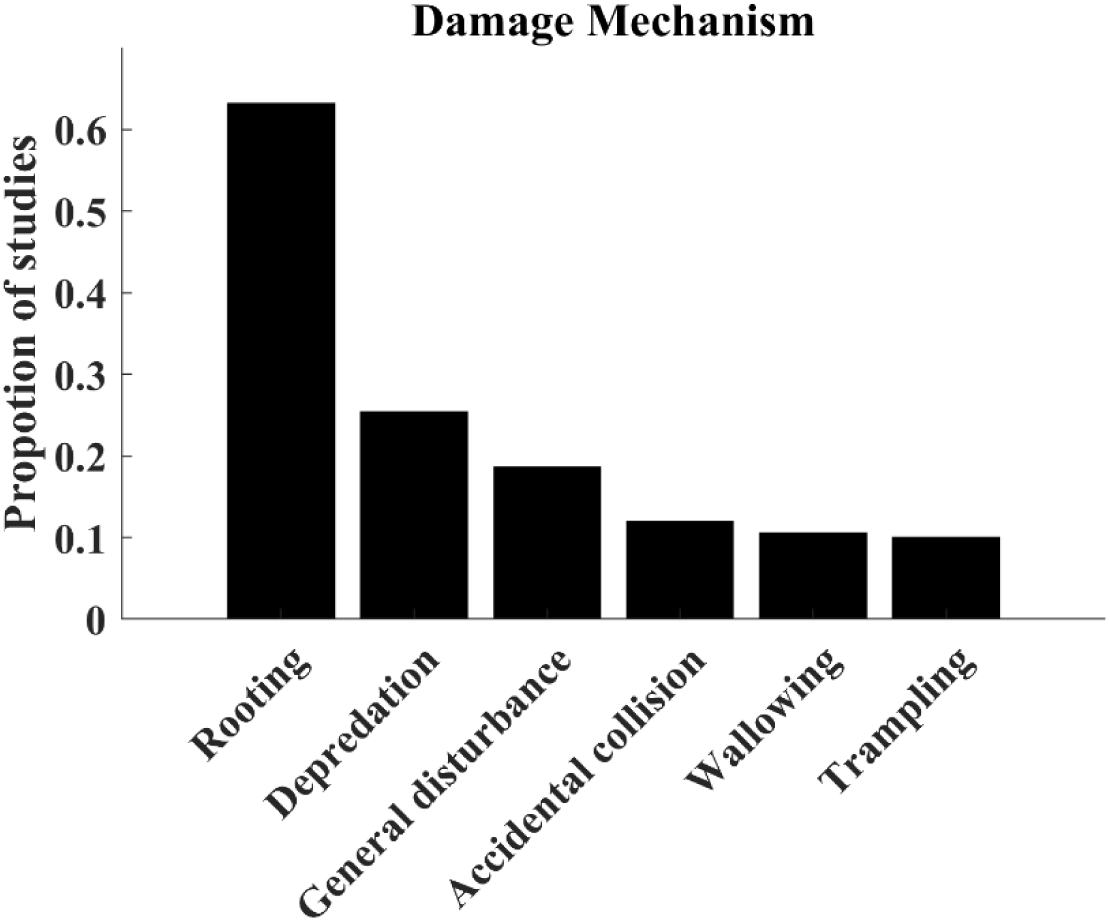
Proportion of studies reporting different mechanisms of damage. Note that these are not mutually exclusive; some studies reported multiple damage mechanisms

These patterns highlight two important insights. First, our understanding of wild pig damage is mainly limited to rooting damage of some natural resources (e.g., wetlands), some crops (e.g., grains, nuts, and beans), and some property (e.g., golf courses and cemeteries), but other damage mechanisms and effects on pasture, livestock, sensitive species, public recreation spaces, and historical sites are infrequent (Figure 3). 12% of the studies we reviewed focused on vehicle collisions, which accounted for > 98% of studies on and human safety.. Thus, the full scope of potential damage caused by wild pigs is unknown for many areas. Second, designing studies to capture the full scope of damage in any local area needs a broad array of techniques to accurately measure each type of damage and its relative importance, because damage can occur to different commodities and natural resources and by different mechanisms.

Damages can also occur from wild pigs simply being in the wrong place at the wrong time, with vehicular collisions. As wild pigs have been expanding and increasing in population, these collisions have become a larger concern because they result in economic and human safety damages (Thurfjell et al. 2015). Of the 25 studies regarding vehicular collisions with wild pigs, ∼19 of those quantitatively focused on identifying patterns in when and where collisions have been reported. The frequency of collisions was influenced by a range of environmental and behavioral parameters including animal densities, time of day, seasonality, distance to habitat cover, and potential hunting pressure (Morelle et al. 2013, Svensson et al. 2014, Kucas and Balciauskas 2020). Mitigating these collisions may include reducing densities of wild pigs, fencing to exclude wild pigs from roadways, construction of roadways away from preferred habitat types, modifications to roads (i.e. reduction in speed limit, lane width, slope and curvature), and modifications to driving behavior (Primi et al. 2009, Diaz-Varela et al. 2011, Beasley et al. 2014, Valero et al. 2015, Thurfjell et al. 2015). However, no studies have evaluated these techniques for reducing damage.

In terms of the amounts of damage that were reported shares of areas damaged ranged from 0.0004 to 0.39 km^2^ of damage per km^2^ with a median value of 0.07. However, only 10.5% of studies measured crop or natural resource damage in terms of area damaged. Monetary values (USD) per km^2^ ranged between $0.40 to $115,680 with a median value of $453.29 of damage per km^2^. However, only 13.4% of studies reported a monetary value for damage. From these findings and similar to the findings of Didero et al. (2023), we recognize that our quantitative understanding of damage is limited relative to our understanding of the presence and type of damage. Also, the wide variety of reported damage measures and different types of damage make it challenging to compare different types of damage in common units, suggesting our understanding of which types of damage cause the most substantial losses remains weak.

## How damage was measured

Our database highlighted methods for measuring damage are variable (Figure 5), even when applied to the same resource. Only 15.3% of studies use field-based methods to measure damage while the rest use indirect damage assessments such as surveys of stakeholders (Figure 6). Roughly 66% of studies did not measure an amount of damage (a gap also noted by Massei and Genov 2004). Roughly 10% of studies provided information for estimating the proportion of damage to a resource. We found that 12.9% of studies measured damage as a number of events (all but 2 studies were vehicle collision studies), while two others (1.0% of all studies) measured weights of crop damaged, and 2.9% measured a number of animals damaged. Only 11.5% of studies reported a monetary value alongside units of area measured. We identified that 23.0% of studies measured damage over time or across multiple locations, but half of these were vehicle collision studies which involve regional analysis. Finally, 12.5% of studies reported a quantitative estimate of wild pig population size (Figure 7, left), but only 4.3% provided a metric for wild pig density alongside damage amounts (Figure 6), and only 2.9% measured damage as an area per unit studied alongside density estimates (i.e., using a comparable metric).

**Figure 5.**
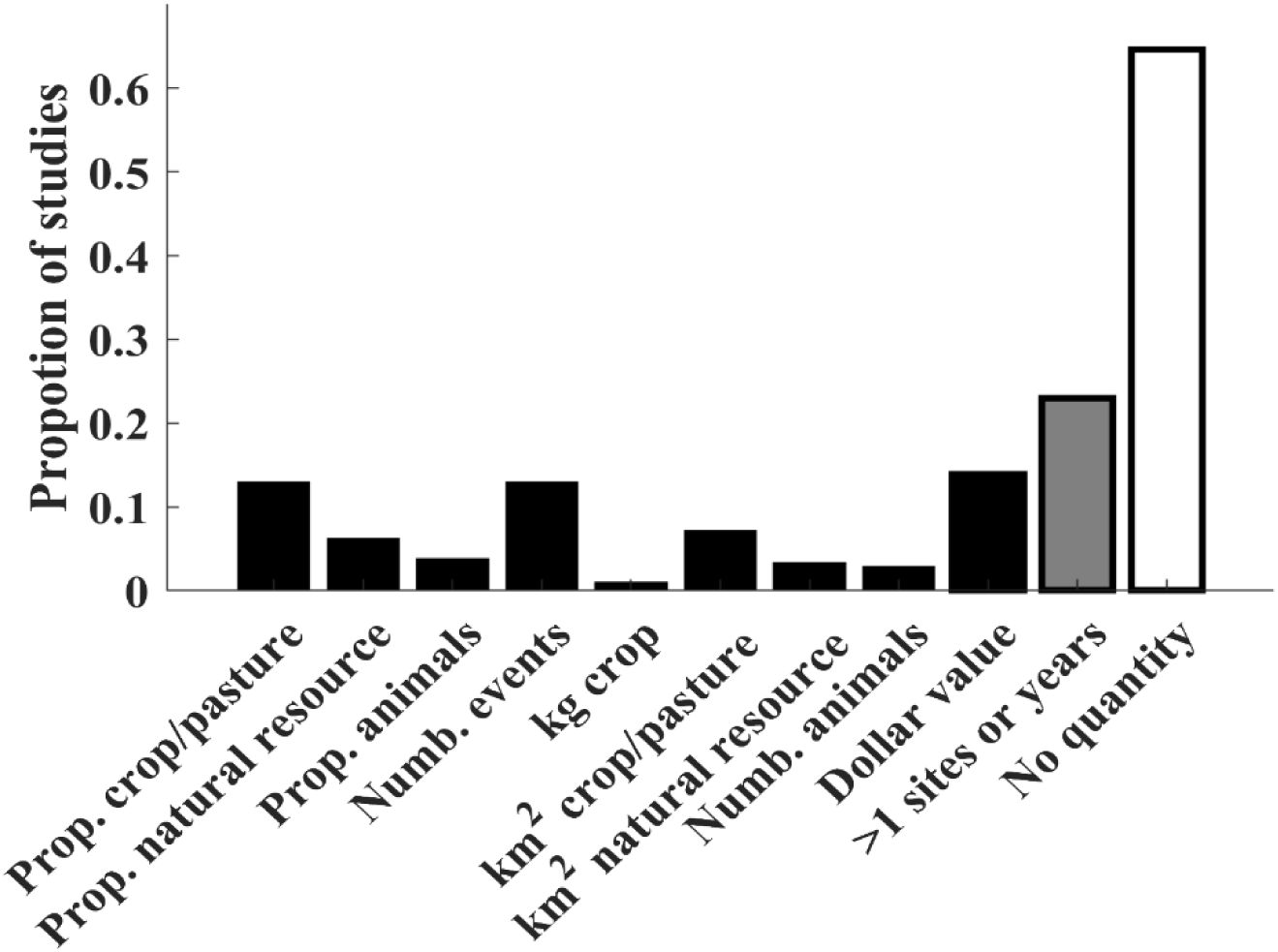
Proportion of studies that report quantitative damage values by different approaches.

**Figure 6.**
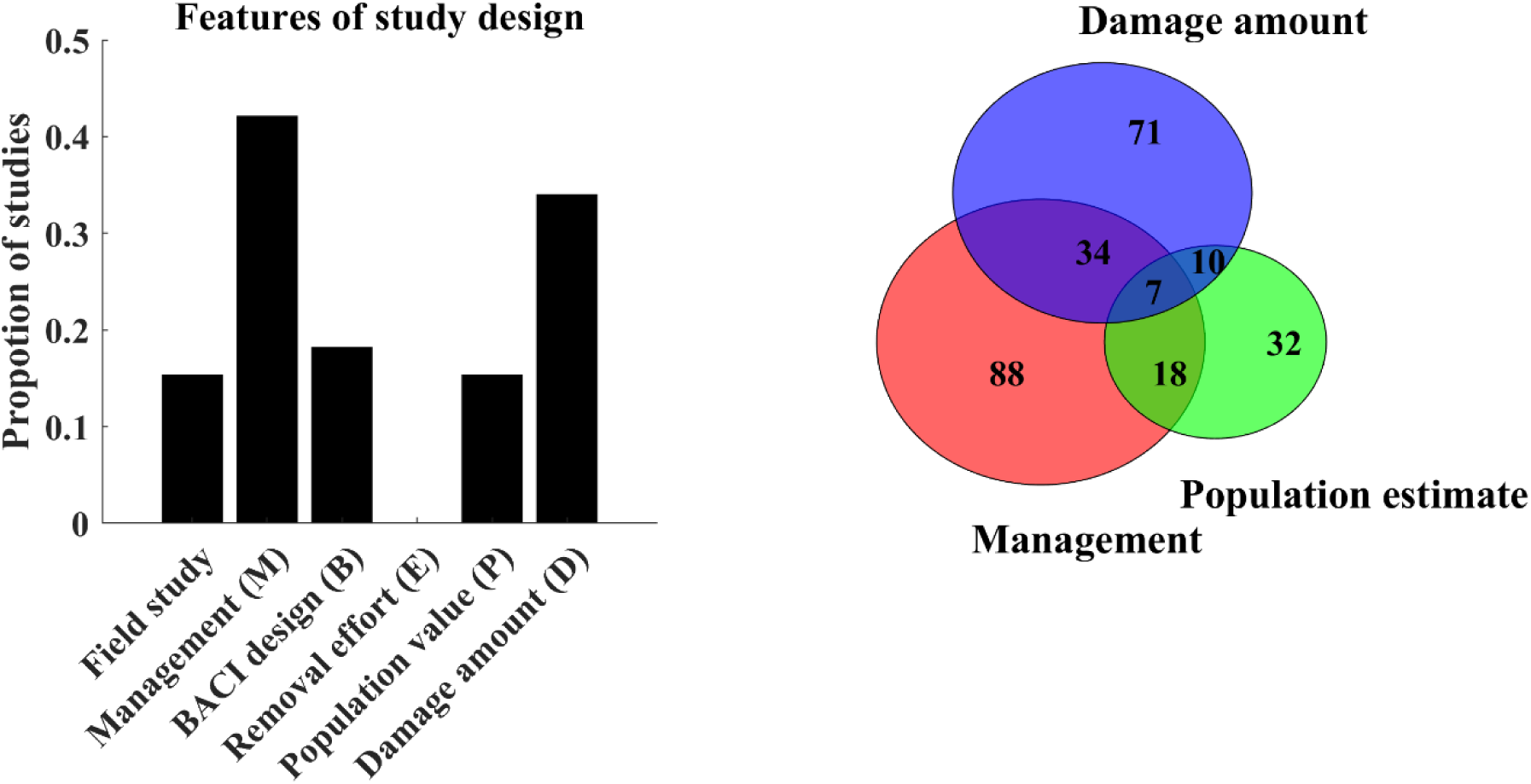
Proportion of damage studies that included different design features (left). Right: Counts of studies that included damage amounts, management, and population estimates. When these design features were combined though, only 3.3% of studies included management, wild pig population estimates, and amounts of damage, 8.6% included management and wild pig population estimates, 16.3% included management and amounts of damage, and 4.8% included population estimates and amounts of damage

**Figure 7.**
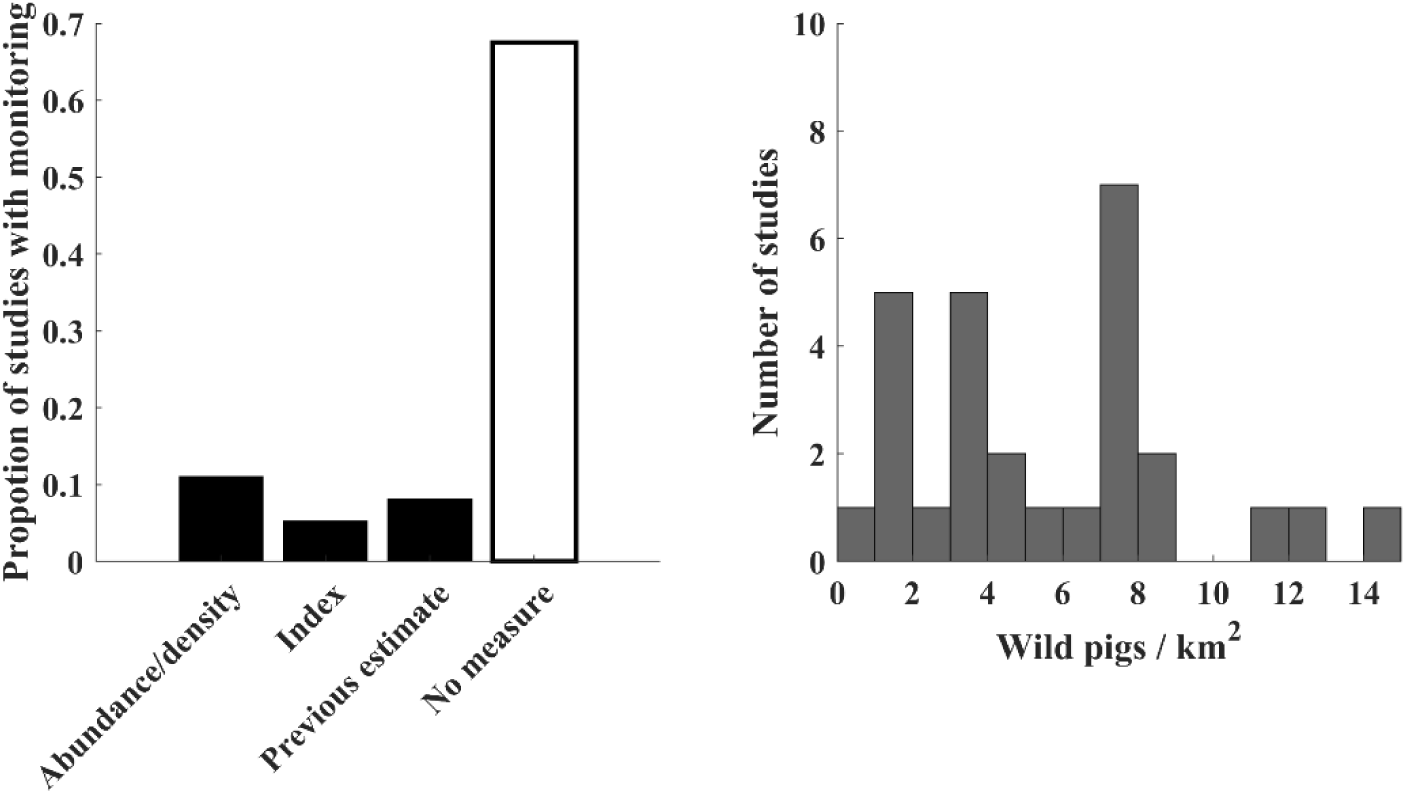
Proportion of 209 studies where an estimate of population abundance was presented. Abundance/density means that abundance or density were measured and estimated as part of the study (12.5% of studies). Index means that an index of abundance or density was estimated as part of the study (6.0% of studies). Previous estimate means that an estimate of abundance, density, or index was reported for the study area but not conducted as part of the current study (9.2% of studies). No measure means that no measure of population abundance or density was reported (7.7% of studies).

In terms of studies that quantified damage amounts, some studies that considered a common damage unit for the numerator (e.g., area of crop damaged), differed in their choice of the denominator. For example, Bobek et al. (2017) measured damages as the percent of the damaged area (all crops) in the potential foraging habitat of wild pigs in farmland, whereas Frackowiak et al. (2013) used the ratio of damage inflicted by wild pigs on hectares of cultivated fields per 1000 hectares of forested area. Alternatively, Mackin (1970) considered the product of the area of damaged crop and the percentage of destruction per 1000 ha of forest. Andrzejewski and Jezierski(1978) reported the average percentage of crops destroyed by wild pigs by type of crop (rye, oats, and potatoes) over the multi-year study period but did not break the estimates out by year. Bleier et al. (2017) focused only on wildlife consumption or injure to corn plants (number of damaged ears vs. total recorded plants) but did not estimate yield losses. Moreover, of the 6 studies that quantified the proportion of crop damaged and wild pig density, methods for estimating metrics of population density were variable. For example, Andrzejewski and Jezierski (1978) reported wild pig density estimates over multiple time points, while Ucarli (2011) estimated average wild pig density over a six-year study period. Bleier et al. (2017) characterized the intensity of space use of wild pigs rather than density. Bobek et al. (2017) and Frackowiak et al. (2013) used the number of individuals harvested per unit area as a proxy of population density, whereas Mackin (1970) used game animal survey data.

## Gaps in damage assessment studies

Two important gaps are that damage assessments for wild pigs have been primarily descriptive or anecdotal (Barrios-Garcia and Ballari 2012) and there is a lack of field-based measures relative to survey-based methods. However, stakeholder reports of damage do not always correlate well with direct measures of damage (Tzilkowski et al. 2002, Humberg et al. 2007), suggesting that reports of total damage within a region are biased. The increasing trend of quantitative damage measures (Table 1) is promising for bridging this gap, but increased field validation of survey-based methods will be important for assessing bias of damage assessments.

Measuring damage over time is another important gap because many types of damage are highly seasonal and thus the presence and impact of damage can change dramatically throughout the year. For example, for a crop such as corn, wild pigs may target the crop only immediately after planting (to consume seeds) or just before harvest (to consume ears of corn; Friesenhahn et al. 2023). Thus it may be important to quantify damage immediately after these times because damage at different stages may be inapparent or have different value to producers, and estimating damage through total crop yield losses could be inaccurate as it could be attributed to multiple causes. Damage to natural resources can also be intermittent, although sometimes with a less predictable seasonal pattern as wild pigs track food resources (Bengsen et al. 2014) or only with short-term effects. Thus a point estimate of the area of rooting damage, for example, may not reflect meaningful damage depending on the ecosystem’s ability to recover. Recent rooting damage might be visible over a large area whereas older rooting damage may have already been replaced by regrowth of vegetation and not visible as rooting damage. However, if past rooting damage causes vegetation shifts that are less favorable for pastures or native species (Bankovich et al. 2016), this is still damage but might require a different technique for measurement and foresight to look for it. Thus, it can be important to measure damage to natural resources both over time and across multiple areas, and to apply different methods to different types of damage to quantify the full scope of damage in terms of how much and how long its effects last.

The amount of effort required to conduct management for reducing damage was not reported alongside damage assessment. Yet, these costs are highly important for valuing the total amount lost due to wild pigs (Carlisle et al. 2021), even though they are not direct measures of damage. Pimentel (2007) estimated that cost associated with damage and management of damage were $1.5 billion (USD) in the USA generated on a per animal basis, but did not take into account differing densities and commodities. In regions with higher densities, the costs of removing wild pigs is substantially lower on a per animal basis (Fischer et al. 2020), which is important to consider.

The density-damage relationship could not be estimated from our database because few studies provided quantitative estimates of wild pig density and damage concurrently (6/209), and those that did used different damage units and density measures. However, our review helped to identify factors to consider when estimating the density-damage relationship. For example, Andrzejewski and Jezierski (1978) and Mackin (1970) found the amount of damage to crops depended on the composition of natural food sources in the nearby environment (an effect that has now been quantified by Wilber et al. (2020) across a wide variety of agro-ecosystems). Other studies showed that crop damage was highest closer to cover (Bleier et al. 2017; Bobek et al. 2017; Frackowiak et al. 2013; Lombardini et al. 2017) and away from urban areas (Lombardini et al. 2017). Thus the relationship between wild pig density and damage is modified by other landscape features that will need to be considered in estimating the relationship between density and damage and using that function to predict damage levels across landscapes more broadly.

A final important gap was monetary valuation of damage to natural resources. Studies that valued damage monetarily focused on valuing damage to crops or private property because that is more straightforward than valuing damage to biodiversity and native habitats. Crops and private property already have a dollar value assigned to them in the human economy, whereas natural resources need to be valued using indirect methods that may require additional data collection (Champ 2017).

## Priorities for damage assessment studies

In Figure 8 we evolve the adaptive management paradigm to inform how decreasing population density correlates to the amount of damage done by the remaining population (Franklin and VerCauteren 2016, Dara 2019). Determining the relationships between wild pig density and damage, and wild pig density and management costs, are essential for evaluating impacts of management programs (Krull et al. 2016; Davis et al. 2018). If these relationships are better understood, the benefits of management can be predicted for an area with a particular profile of resources and wild pig density, even in areas where damage assessments are not available. Damage inference at such a landscape scale improves management planning and prioritization of control resources across a range of already invaded or potentially at-risk areas. Though assessing wild pig densities or abundance can be time consuming and expensive, recognize that a variety of methods (from coarse trend data to hard-earned rigorous density assessments) are employed annually for the purpose of setting harvest quotas for big game species in local areas throughout several developed and undeveloped countries. Further, with rapid advancements in passive sampling tools (e.g., trail cameras, acoustic recording) and analytical methods (Moeller et al. 2018, Gilbert et al. 2020), assessing populations before and after management actions is more practical now than ever before (Davis et al. 2018, Davis et al. 2022).

**Figure 8.**
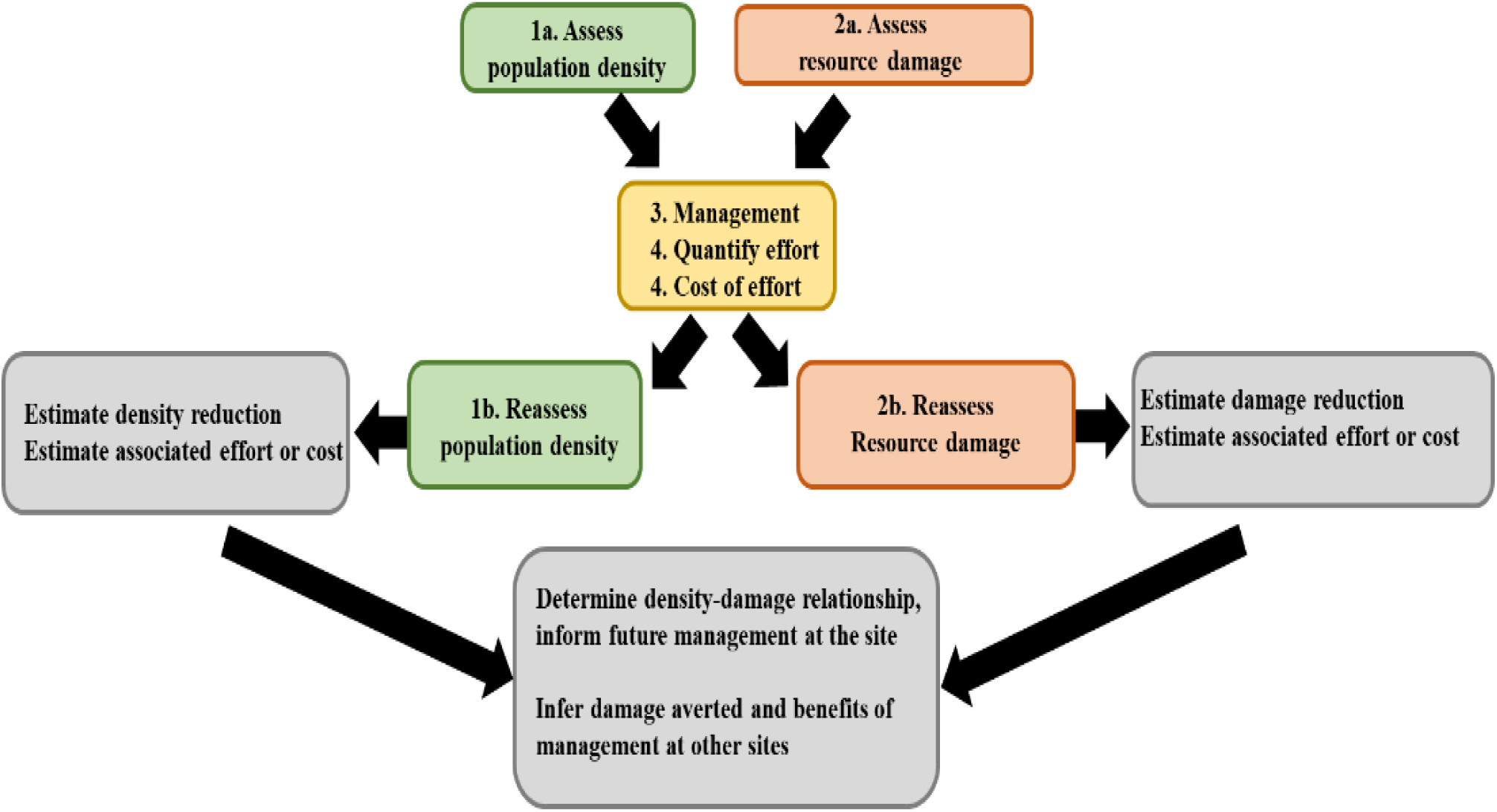
Adaptive framework for estimating management impacts on damage caused by wild pigs. Steps 1a and 2a assess population density (e.g., trend indices, wild pigs/km^2^) and amount of damage (e.g., trend indices, percent or value of resource impacted; standardization across areas allows for comparisons). Steps 3 and 4: Conduct management with intent of reducing damage and document effort and cost. Steps 1b and 2b: Reassess density and damage and quantify reduction relative to effort. Measuring density and damage concurrently at multiple study sites or over time at a site allows estimation of the density-damage relationship. The density-damage relationship allows inference of the amount of damage averted in other areas based on the resources and other landscape features in those areas. Knowledge of the relationship between density and management effort together with an estimate of the density-damage relationship allows inference of total benefits of applying a given amount of management in an area based on damage reduction, damage aversion, and management costs

With the large potential for wild pigs to affect native species, biodiversity, and sensitive ecosystems (McClure et al. 2018), valuing damage to natural resources is high priority to direct the most urgent management actions (Didero 2023). For agriculturally damaging species such as wild pigs, it would be useful if methods for valuing damage to natural resources are developed in a way that these damages can be summed with damages to agriculture for producing total damages based on all locally available resources, allowing evaluation of management benefits and improved prioritization of management actions. A major pressing challenge that remains to be addressed is how to integrate multiple types of damage quantities into an overall value of damage.

Damage assessment studies to specific resources are time-consuming and expensive. Thus for species that cause damage to many different resources, damage assessments tend to focus only on a subset of resources. A framework for guiding damage assessment studies (Figure 8) is important for guiding independent studies to provide inference of total damages. Damage assessment studies should aim to report quantities where possible, and in a manner that they can be standardized within a particular type of damage. For example, reporting 1000 km^2^ of crop damage without reporting the amount of area that was surveyed for crop damage or the time frame that the damage occurred in makes it difficult to compare the significance of the damage amount across studies. This is especially important for species that cause damage to many different resources (e.g., wild pigs) because damage assessment to different sectors is often conducted by researchers from multiple disciplines using different techniques. Damage amounts should be described as proportions of area/amounts per unit time to different resources (with the denomitors for space/amount and time reported – i.e., damage rates) so that individual elements can be compared on equal grounds and total damage can be compiled.

Most damage assessments have focused on quantifying current damage. However, preventing damage to new areas is often a goal of management programs (Hygnstrom et al. 2014). Quantifying potential damage alongside current damage can help with planning optimal control strategies and evaluating program benefits more comprehensively. Potential damage can be estimated by linking estimates of the amount of damage that wild pigs cause to predictions of where wild pigs will spread (Snow et al. 2017) and which resources occur in those areas. Estimates of management impacts on preventing the spread of wild pigs to particular areas (Pepin et al. 2019) can be combined with data on the amount of at-risk resources in those areas to predict the amount that would likely be damaged if management does not occur.

## Conclusions

Knowledge of damage caused by invasive species drives the inception, funding, and planning of management programs. Our survey of 209 damage assessment studies for wild pigs revealed highly variable damage assessment methodologies and objectives, even when measuring damage to the same resource. Measuring damage amounts and standardizing those measures per unit area or amount per unit time is an important goal for future studies. This will allow collation of damage estimates from multiple studies to provide more comprensive estimates of damages and costs caused by wild pigs. Also, currently there is a lack of wild pig density or indices of population change information reported alongside damage estimates and control efforts, and thus little knowledge of the relationship between abundance, damage, and management impacts. Estimates of these relationships and how it varies are needed for understanding the level of management that is needed (i.e., population control) based on the local conditions (i.e., population abundance of wild pigs) and the severity of damage that is ensuing (i.e., crop depredation). Overall, we recommend the following priorities for future studies of damage from wild pigs to strive for: 1) report the amount of damages in a standardized fashion (e.g., area damaged/area surveyed), 2) evaluate and report the amount of damage relative to the density of wild pigs, and 3) when reporting economic costs of damages incurred and management actions, describe the economic valuation method used along with the year of reference for the valuation. Capturing these elements are necessary steps to predict the benefits of management for an area with particular profile of resources and wild pig density, even in areas where damage assessments are not available, allowing for more generalizable results that can inform managers across the nearly global distribution of wild pigs.

## Acknowledgements

The authors thank Michael Lavelle and Joseph Halseth for helpful input. Funding for this research was from the United States Department of Agriculture, Animal and Plant Health Inspection Service (APHIS), Wildlife Services, and the APHIS National Feral Swine Damage Management Program.

## Statements and Declarations

Funding for this research was from the United States Department of Agriculture, Animal and Plant Health Inspection Service (APHIS), Wildlife Services, and the APHIS National Feral Swine Damage Management Program.

The authors have no relevant financial or non-financial interests to disclose.

All authors contributed equally in developing the manuscript.

## Notes

### Competing Interest Statement

The authors have declared no competing interest.

